# Linearization Autoencoder: an autoencoder-based regression model with latent space linearization

**DOI:** 10.1101/2022.06.06.494917

**Authors:** Sangyeon Lee, Hanjin Kim, Doheon Lee

## Abstract

Regression analysis is one of the most widely applied methods in many fields including bio-medical study. Dimensionality reduction is also widely used for data preprocessing and feature selection analysis, to extract high-impact features from the predictions. As the complexity of both data and prediction models increases, it becomes important and difficult to interpret the model. We suggested a novel method, linearizing autoencoder, for regression analysis with high-dimensional data. Based on the autoencoder model, we introduce a novel loss function to make data points aggregate corresponding to their known labels and align them preserving linear relations of the known feature. This model can align data points to the linear relations of labels, and achieve both the prediction and feature selection performances by extracting features that are important to the label we want to predict. Also, we applied this method to the real-world data and the result indicates that this method can successfully disentangle the latent space with given centroids in a supervised manner. This method can be applied to various prediction problems in biomedical fields.

## INTRODUCTION

Regression analysis is to predict continuous variables with given data. It is widely used in various fields of science and technology, especially in the bio-medical field, predicting gene expression patterns [1, 2, 3, 4], biomedical signal analysis such as electrocardiography [5, 7] and electromyography [6], disease diagnosis from patient data [8, 9, 10], drug dose prediction [11, 12, 13, 14], and many other fields. Basically, linear regression is widely used in these kinds of prediction tasks [1, 2, 5], but as the data complexity increases, advanced regression methods including regression trees such as extreme gradient boosting (XGBoost) are suggested and vividly applied [14]. Moreover, the fast development of deep-learning techniques makes artificial neural network-based prediction models suitable for analyzing high-dimensional and unstructured complicated data [8, 9], and they tend to outperform compared to classical machine learning algorithms.

However, the more complex the model, the more difficult it becomes to interpret the model. In order to interpret complex models, there are tries to extract hidden features which has a high impact on the prediction model. One of those tries, dimensionality reduction is a widely used concept to analyze high-dimensional data in various fields. [15, 16, 17] Dimensionality reduction aims to represent high-dimensional data to low-dimensional space while preserving some meaningful features of the original data. The interpretability of the machine learning model can be improved by removing noises and multicollinearities through dimensionality reduction. Traditionally, linear methods [18] such as principal component analysis (PCA) [19, 20] and singular vector decomposition (SVD) are widely used, [21, 22] and to handle non-linear features of the data, non-linear methods like manifold learning are suggested [23, 24]. Also, with the development of deep neural network techniques, autoencoders became one of the most used methods for dimensionality reduction. As above-mentioned methods are successfully projecting high-dimensional data into low-dimensional latent space preserving hidden properties of data, which is shown by good task performance [25, 26]. But still, interpretability of latent space remains an important factor because intrinsic information hidden in high-dimension is implicated in the latent space. Studies like latent space disentanglement are trying to connect features in the latent space to observable features in high-dimensional space for improving latent space interpretability and overall downstream task’s general performance. [27]

While these methods can disentangle the latent space and find important features that distinguish the data, for general regression problems, it is important to find features that are highly related to the value we want to predict. As a result, a feature selection method to connect features represented in the latent space to observable high-dimensional features which are important to the predictions which is a continuous value. Here, we developed a modified autoencoder model which can be applied to general regression tasks. With the modified loss functions, this linearizing autoencoder model can project data to low-dimensional space considering the linear relations of the value we want to predict. This model achieves both high performances for the regression task and disentangling latent space, and shows the possibility to be applied in various fields.

## RESULT

### MODEL ARCHITECTURE

Linearizing autoencoder is based on autoencoder combining encoder and decoder consists with several fully-connected hidden layers. (Fig 1, a) We adjusted the loss function of the autoencoder to achieve both reconstruction of the given data, the prediction performance of the given task, and interpretable linearized latent space. We combined four loss functions, mean squared error (MSE), average intraclass distance, average interclass distance, and the average distance from the label-wise centroid to achieve the aforementioned purpose. (Fig 1, b) First, the MSE measure of the difference between original data and reconstructed data is one of the generally used loss functions in autoencoders to improve the performance of data reconstruction. For *n* data represented as a vector *X*, MSE can be calculated by:

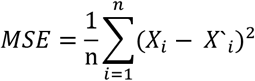

**Figure 1.**
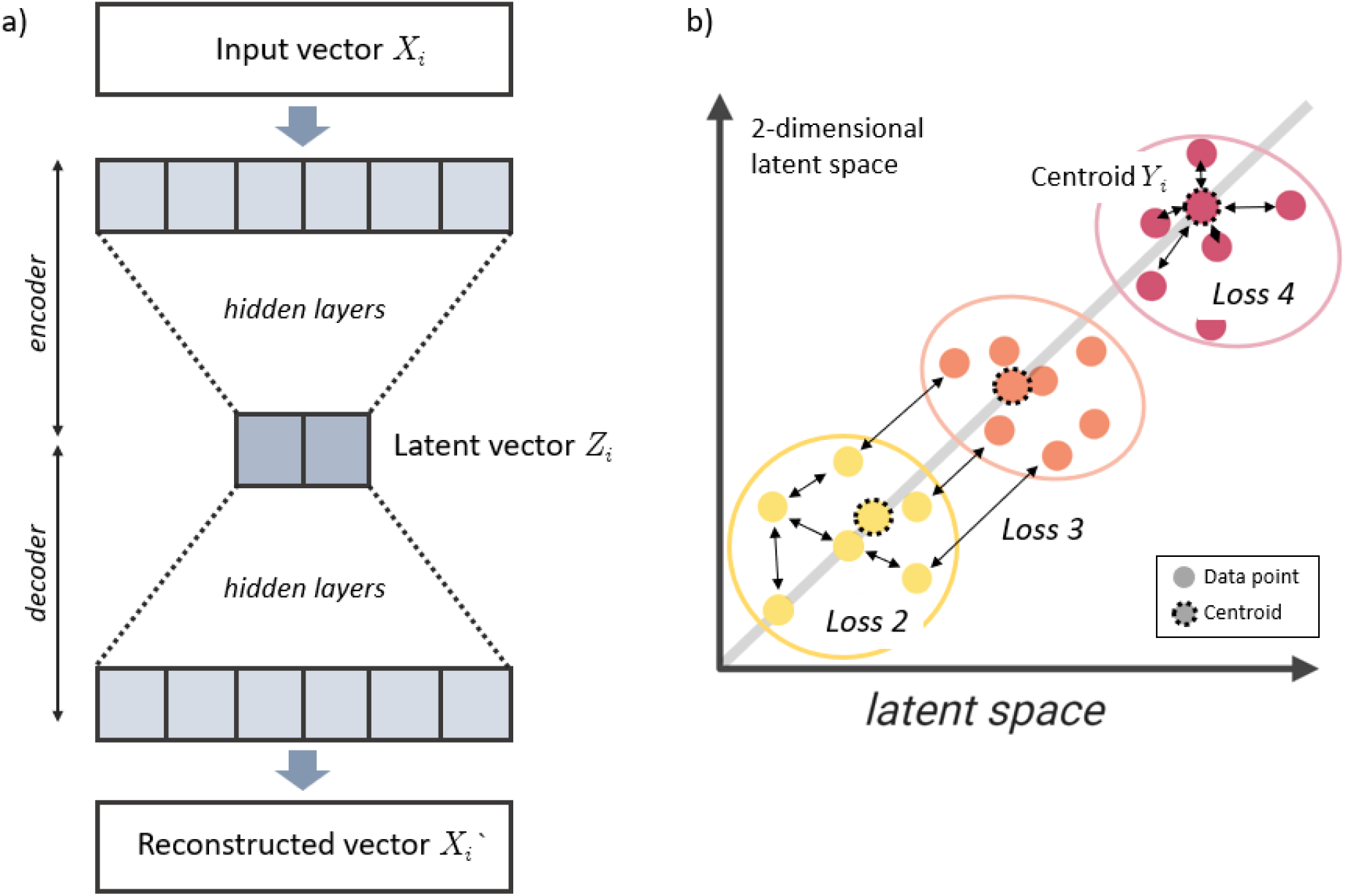
Simple schematic of linearizing autoencoder and its loss function. (a) The structure follows general form of autoencoder model which combines encoder and decoder. It takes a vector as an input, project it to 2-dimensional latent space through hidden layers, and reconstructs a vector which is similar to the input vector. (b) The global loss function consists of four loss functions. Loss 1, which is not in the figure calculates the difference between the input vector X and reconstructed vector X’. Loss 2 measures the average distance of data of the same class in the latent space. Loss 3 calculates average distance of data of the different class in the latent space. For loss 4, the model introduces the centroid for each class with the known value for prediction, and calculates the average distance of data in the class between the centroid.

For good prediction performance and the linearized latent space tuned by an interesting label, data points of the same label should be aggregated and those of the different labels should be separated in the latent space. For this purpose, we added averaged intraclass distance and averaged interclass distance as loss 2 and loss 3 respectively. These loss functions improve the performance clustering by placing data points with the same label close and data points with a different label distant by minimizing the averaged intraclass distance and maximizing the averaged interclass distance. Because the neural network model is trained to minimize the loss functions, the averaged interclass distance is inverted before calculating the global loss. Loss 2 and loss 3 are calculated as:

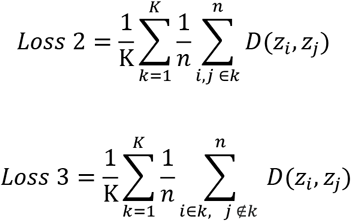

where *K* is the number of given labels, *n* is the number of the data which belongs to the current label *k, z* is the coordinate of the data points in the latent space, and *D* denotes the distance function. Here we used Euclidean distance for the calculation.

Additionally, we added loss 4 to the loss function to reflect the known labels of the dataset and to linearize the latent space. We arbitrarily set the centroids for each label on the latent space. These centroids represent the ideal location of data points that correspond to the known label and are used to gather those data points. To make the data point closely located to the centroid of the label, we calculated the averaged distance between the data point and the centroid in the latent space. By minimizing the distance, data points will aggregate to the centroid, keeping the linear feature of the data. Also, centroids can disentangle the latent space and make the latent space and prediction model interpretable because their coordinates on the latent space are determined with previous knowledge on the nature of label value. For vector *X*_*i*_ with label *y*_*i*_, loss 4 in the latent space is calculated as:

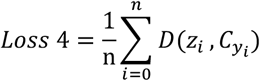

where the *C*_*yi*_ is the centroid for given label and in the 2-dimensional space. The centroid is determined as (*y*_*i*_, *y*_*i*_) in this study. As a result, the total loss function in this method is defined as:

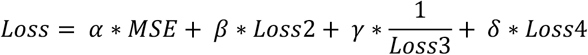

where *α, β, γ* and *δ* are weight coefficients for each loss value, which are determined heuristically.

### APPLICATION TO REAL DATASET

To test the linearizing autoencoder, we applied this model to a drug dose prediction task [14] as a benchmark. The data provides demographic, drug combination, and laboratory information of 184 patients who are administrated vancomycin. 26 features and a therapeutic dose of vancomycin (500, 1000, 1500, 2000, 3000 mg/d) are provided for a patient. 26 features of a patient are directly used as an input vector and the dose of vancomycin is used as a label to predict. We trained the linearizing autoencoder using 100 patients as training data. To align each data point in the latent space in order of therapeutic dose, we assigned label-wise centroid with dose information. Hyperparameters including the structure of hidden layers, batch size, learning rate, weight decay, and weight coefficient of each loss function are heuristically determined. During the training, the total loss reduced as four losses reduces, as shown in figure 4 (a), showing that four loss functions we applied in this task works well.

After the training, we validated our model using the rest of the patient data. We compared the latent space of three models, an ordinary autoencoder that uses only MSE for loss function, an autoencoder for clustering tasks with MSE, averaged intraclass distance, and inverse of averaged interclass distance, and linearizing autoencoder which combines averaged distance between class-wise centroids and aforementioned loss functions. (Fig2 a, b, c) As shown in figure2 a, b, and c, combining MSE and averaged intraclass and interclass distance to a loss function successfully separates data points in different labels, but not orders of labels. (fig2, c) It can classify a given data to known labels, but cannot predict actual values between or above known labels. However, by linearizing the latent space by minimizing the averaged distance between data points and the centroid, actual values that are not covered in the training data can be predicted like a linear regression model. (fig2, d)

**Figure 2.**
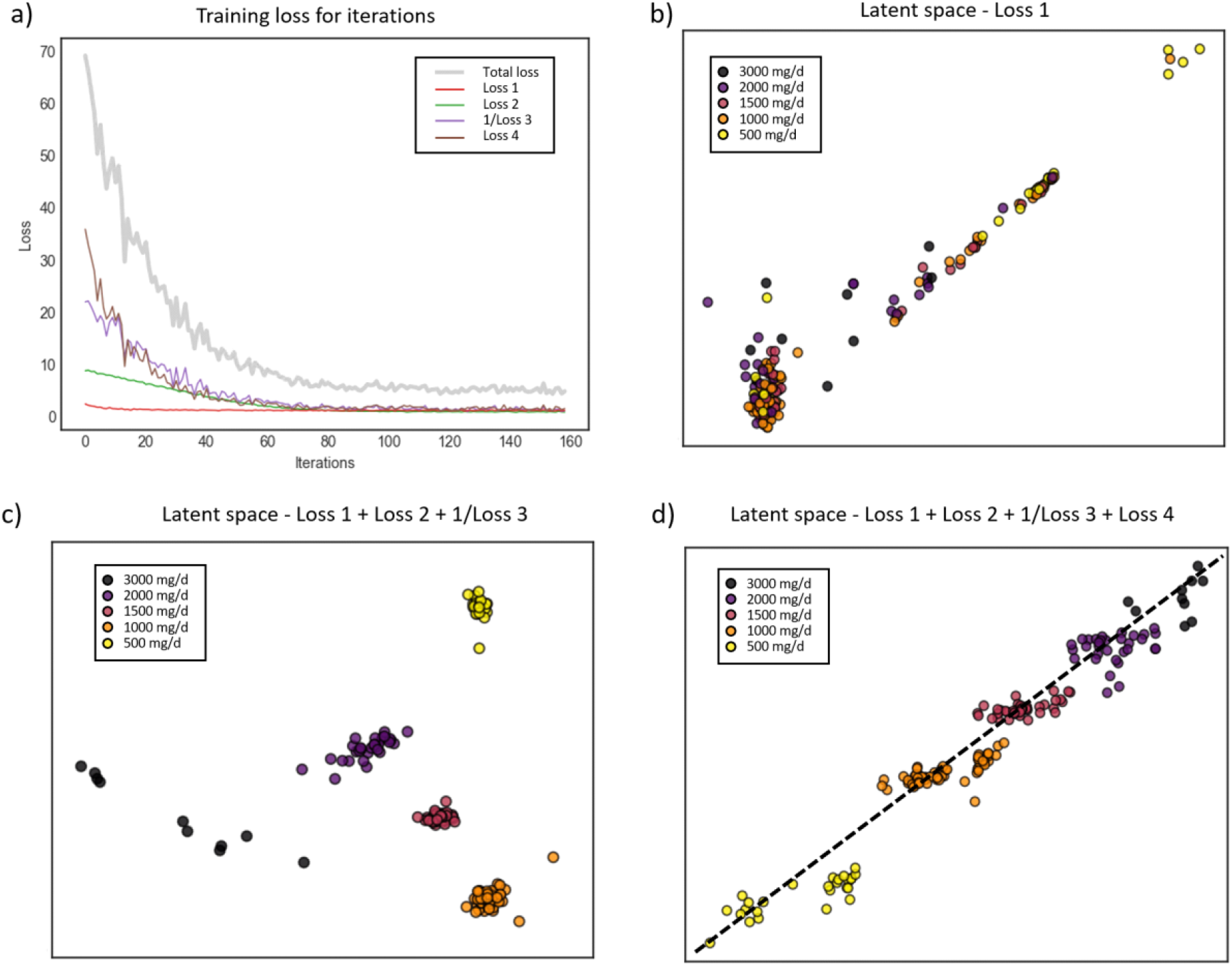
Example of linearizing autoencoder with drug dose prediction task. a) The training loss reduces as the training progresses. b) The distributions of data points in the latent space after the training with only MSE for the loss function. c) The latent space after the training with the loss function containing MSE, the averaged intraclass distance, and the averaged interclass distance. d) The linearized latent space with the drug dose value, trained by the loss function that combines all aforementioned loss functions and the averaged distance to the label-wise centroids.

## METHOD

### Drug dose prediction

As a neural network-based model, there are many hyperparameters to be tuned. We manually set the batch size to 32 and followed some widely-accepted values for learning rate (=1e-4), and weight decay (=1e-4), and heuristically determined the other hyperparameters such as the structure of the autoencoder, and the number of training epochs. We set five centroids to calculate loss 4 during the training. Labels are normalized from 0 to 1, and centroids are determined corresponding normalized value of the label. For example, the maximum and minimum labels are 3000mg/d and 500mg/d, so normalized labels will be 1 and 0 respectively. Centroids are placed in the line y=x, so the coordinate of centroids in the latent space is (normalized label, normalized label), in the case of labels 500mg/d and 1000mg/d, (0, 0) and (0.2, 0.2) will be the centroids respectively.

## CONCLUSION

We have proposed a linearizing autoencoder model, which can be applied to general regression tasks. Based on the autoencoder model, it learns how to reduce the dimensionality of given data. While training, it learns to linearize the latent space by aligning data points to the value we want to predict. For this purpose, we suggested a new loss function, combining MSE, averaged intraclass distance, averaged interclass distance, and averaged distance between class-wise centroids. The proposed method was applied to the benchmark data, which aims to predict drug dose from patient data. The result indicates that the proposed method successfully aligns data points corresponding to their known labels compared to autoencoder models with different loss functions, which means it learned linear relationships of data points. These data points in the latent space can be directly used for prediction tasks or classification tasks and to further feature selection tasks. It also shows the possibility to be applied general regression tasks.

## Notes

### Competing Interest Statement

The authors have declared no competing interest.

